# Short- and mid-term temporal variability of the human urinary microbiota: an observational cohort study

**DOI:** 10.1101/2024.06.25.600589

**Authors:** Vojtěch Tláskal, Jan Hrbáček, Vítězslav Hanáček, Petra Baránková, Pavel Čermák, Roman Zachoval, Priscila Thiago Dobbler

## Abstract

Understanding the temporal variability of the microbiome is critical for translating associations of the microbiome with health and disease into clinical practice. The aim of this study is to assess the extent of temporal variability of the human urinary microbiota.

A pair of catheterized or mid-stream urine samples were collected from study participants at 3–40-month interval. DNA was extracted and the bacterial V4 hypervariable region of the 16S rRNA gene was sequenced on the Illumina MiSeq platform. The alpha and beta diversity of paired samples was analyzed using Chao1 and Shannon indices and PERMANOVA.

A total of 63 participants (43 men and 20 women with a mean age of 63.0 and 57.1 years, respectively) were included in the final analysis. An average of 152 ± 128 bacterial operational taxonomic units (OTUs) were identified in each urine sample from the entire cohort. There was an average of 41 ± 32 overlapping OTUs in each sample pair, accounting for 66.3 ± 29.4% of the relative abundance. There was a clear correlation between the number of overlapping OTUs and the relative abundance covered. The difference in Chao1 index between paired samples was statistically significant; the difference in Shannon index was not. Beta diversity did not differ significantly within the paired samples. Neither age nor sex of the participants influenced the variation in community composition. With a longer interval between the collections, the relative abundance covered by the overlapping OTUs changed significantly but not the number of OTUs.

The abundance of bacterial taxa present in both collections fluctuated, but the proportion of these taxa in the community was about two-thirds of the total bacterial community. No significant intraindividual differences in beta diversity were observed between the two urine samples.

**Data Summary:** The raw DNA sequences together with anonymized sample metadata have been deposited at the NCBI SRA under the accession number PRJNA1093489. Processing scripts are deposited at a public repository https://doi.org/10.5281/zenodo.12556460. Processed sequencing files and tables including full taxonomic assignment are deposited at https://doi.org/10.6084/m9.figshare.26046355.

**Importance:** This study represents a comprehensive investigation dedicated specifically to the subject of urinary microbiome (UM) temporal stability. Our dataset (n = 63) consists of a relatively large group of patients of one ethnic origin but varying ages and both sexes. Additionally, samples from individual participants are separated by different lengths of time. This approach allows us to assess the effects of three variables on the stability of human UM. Our findings demonstrate that, while the relative abundance of dominant bacteria varies, repeated collections generally share more than 60% of the bacterial community. Furthermore, we observe little variation in the alpha and beta diversity of the microbial community in human urine. These results help to understand the dynamics of human UM and enable interpretation of future studies.

## Introduction

The existence of the human urinary microbiota (UM) is now considered a fact. (1,2) Both the lower and upper urinary tract harbor diverse microbial communities, some members of which have been cultured using expanded quantitative urine culture, while others have been detected by next-generation sequencing of the 16S rRNA gene. There is evidence that, in addition to the bacteriome, the UM also includes viruses (the virome) (3)and fungi (the mycobiome)(4), about which very little is known to date.

As far as the niches of the human body are concerned that have been incorporated in the Human Microbiome Project (5), their microbial communities appear to be stable. In the gastrointestinal tract, the composition of the microbiome is generally consistent over time and even after antibiotic exposure or a temporary change in diet, the microbiota returns to its original state. Not only the operational taxonomic units (OTUs) but also the metagenomic profile remain stable within a given individual. (6) Despite the exposure of the skin to various environmental factors, the stability of the community is maintained in the short and long term. A prolonged interval between sampling did not lead to an increase in the variation of the cutaneous microbiota within an individual. (7) In the oral cavity, the subgingival microbiota remains stable to a certain extent, although it varies from person to person and even between different sites of the oral cavity. (8) The stability of the vaginal microbiome over time is more challenging to determine due to the physiologically ever-changing influences of the hormonal cycle, perturbations of the microbiota during pregnancy, etc. (9)

Understanding the temporal variability/stability of the microbiota is crucial for translating associations of the microbiome with health and disease into clinical practice. However, until now it has been unclear how stable the UM is or how reliably a single sample reflects the microbial communities of a bladder over time. The present study aims to fill this knowledge gap and assess the degree of temporal variability of the human UM.

## Methods

### Population

The population of the present study is based on a cohort of patients with bladder cancer and a control group of subjects without cancer reported previously (10). A total of 107 subjects were asked to provide a second urine sample for the assessment of their urinary microbiota stability. Those willing to re-attend provided a second sample of urine for the assessment of alpha and beta diversity. To expand the study cohort, members of the research staff (healthy volunteers) were asked to provide two samples of urine several months apart. Antibiotic treatment was not administered to any subject six weeks prior to the first urine collection. The study was conducted in accordance with the Declaration of Helsinki after previous approval by the Ethics Committee of the Institute for Clinical and Experimental Medicine and Thomayer Hospital with Multi-center Competence under the number G-19-01 and informed consent was obtained from all participants prior to enrolment. Participants’
s enrollment into the study and sample collection was conducted between 27 May 2019 and 31 January 2023. The STORMS checklist for this study is attached as S1 Table. (11)

### Sample collection and handling

The first paired sample was collected either as a mid-stream, clean-catch voided specimen of urine (mid-stream urine, MSU) or obtained via transurethral catheterization under anesthesia before a surgical procedure. The second paired sample was uniformly collected as MSU. An aliquot of each sample was stored at 4°C and processed within 24 hours with standard urine culture; another aliquot was frozen at -20°C on the same day for several months until shipping, DNA extraction and sequencing.

### DNA extraction and sequencing

DNA extraction and 16S rRNA gene sequencing were performed as previously described (12) at the Institute of Microbiology of the Czech Academy of Sciences, Prague, Czech Republic. Briefly, DNA was extracted from urine samples using Eligene Urine Isolation Kit (Elisabeth Pharmacon, Ref. 90051-50) according to manufacturer’s instructions in a random order of patient groups to avoid batch effects. The whole amount of urine was vortexed for 15 s, 10 mL of urine was then centrifuged at 6,000× g for 20 min, the supernatant was discarded, and pellet resuspended in 200 μL of molecular grade water, 200 μL of MI3 solution, and 20 μL of Proteinase K was added. After 15 s vortexing, the mixture was incubated for 15 min at 65 °C. The lysate was centrifuged at 6,000× g for 5 min. The supernatant was transferred to microtube and 210 μL of MI4 solution added. The lysate was centrifuged for 1 min at 13,000× g. Due to a low microbial load of source samples, DNA extraction controls as well as negative controls for PCR reactions were included. Enrichment of microbial DNA was not performed. Yield and purity of extracted DNA were checked using NanoDrop 1000 Spectrophotometer (Thermo Fisher Scientific). Samples with even as low DNA concentration as 5 ng μL^-1^ were included in the amplification step.

The primers 515F (5’-GTGCCAGCMGCCGCGGTAA) and 806R (5’-GGACTACHVGGGTWTCTAAT)(13) were used to amplify the hypervariable region V4 of the 16S rRNA gene. Each forward primer was barcoded by a custom sequence of nucleotides designed to multiplexing of different samples. PCR was performed in triplicates, and every reaction contained 5 μL of 5× Q5 Reaction Buffer for Q5 High-Fidelity DNA polymerase (New England Biolabs, Ref. B9027S); 0.25 μL Q5 High-Fidelity DNA polymerase (New England Biolabs, Ref. M0491L); 5 μL of 5× Q5 High GC Enhancer (New England Biolabs, Ref. M0491L); 1.5 μL of BSA (10 mg mL^-1^, GeneOn, Ref. 209-005W); 0.5 μL of PCR Nucleotide Mix (10 mM, Thermo Fisher Scientific, Ref. R0191); 1 μL of primer 515F (10 μM); 1 μL of primer 806R (10 μM,); 1.0 μL of template DNA and sterile ddH_2_O up to 25 μL. Conditions for amplification started at 94 °C for 4 min followed by 25 cycles of 94 °C for 45 s, 50 °C for 60 s, 72 °C for 75 s and finished with a final setting of 72 °C for 10 min.

The successful PCR was confirmed using agarose gel electrophoresis. Three PCR reactions were pooled together in order to randomize PCR bias and to get higher DNA yield. Pooled samples were purified by MinElute PCR Purification Kit (Qiagen, Ref. 28004) and mixed in an equimolar amount according to the concentration measured on the Qubit 2.0 Fluorometer (Thermo Fisher Scientific). Sequencing libraries were prepared using the TruSeq DNA PCR-Free Kit (Illumina, Ref. 20015962) according to manufacturer’s instructions while following official ligation protocol for the kit used. Sequencing was performed on Illumina MiSeq in a 2 × 250 bases sequencing run. Raw fastq files were retrieved and used as input to the bioinformatic pipeline described in the following section.

### Statistical analyses

Demographic and clinical data were analyzed as continuous or categorical variables and reported as mean and standard deviation (SD)/interquartile range (IQR) or counts with percentages as appropriate.

The sequencing data were processed using SEED 2.1.2 (14). Pair-end reads were merged using fastq-join (15). Sequences with ambiguous bases were omitted as well as sequences with average quality PHRED score <30. The chimeric sequences were detected and removed using USEARCH 8.1.1861, and clustered into OTUs using the UPARSE algorithm (16) at a 97% similarity level. This step yielded in average 15,107 ± 8,700 sequences per sample (SD, min = 129). To filter out non-bacterial sequences, the most abundant sequence from each OTU was assigned to the closest hit from the GenBank database by NCBI BLAST 2.10.1. Taxonomy was assigned using DECIPHER 2.30.0 (17) and threshold 40 with IDTAXA algorithm trained on SILVA SSU database r138 (18). This step yielded in average 11,524 ± 8,669 sequences per sample (SD, min = 19).

If one of the paired samples showed Goods sequence coverage <85, the pair was excluded from analysis. Global singleton OTUs were removed before analysis of OTUs occurrence and relative abundance of individual OTUs was calculated. Alpha and beta diversity analyses were performed using the packages tidyverse 2.0.0 (19), vegan 2.6-4 (20), QsRutils 0.1.5 (21), GUniFrac 1.8 (22) and tidylog 1.0.2 (23) in R 4.3.1 (24). For alpha diversity analyses, rarefaction was done following (25): each sample was randomly subsampled to 1,000 sequences while excluding patients with at least one sample below this threshold (S2 Table). Subsampling was iterated 1,000 times and per-sample average summary statistics was calculated. The following indices were calculated (26): Chao1 (reflecting species richness (22) and Shannon diversity index (reflecting species evenness). Normality was tested by Shapiro–Wilk test, differences among groups were tested by Kruskal-Wallis and Wilcoxon paired test as appropriate and PERMANOVA based on the Bray–Curtis dissimilarity matrix. Temporal trends in the number of shared OTUs and their relative abundance were modelled by linear regression. The results were considered statistically significant at the level p < 0.05.

## Results

After exclusion of samples with insufficient Good’
ss coverage (6 subjects), a total of 63 subjects and their paired samples of urine were included in the analysis (S2 Table). These samples were provided by 43 men (mean age 63.0 ± standard deviation (SD) 15.2 years) and 20 women (mean age 57.1 ± 13.6 years). The time gap between the two collections ranged from three to 40 months (median 24.8, IQR 14.9 months).

Standard urine culture was negative at 10^5^ colony forming units/mL for all samples except one (no. 307) at the first sampling. The second sample was culture-negative in 61 of subjects. Among the study participants, 28 were patients with non-muscle invasive bladder cancer (stage Ta-T1) presenting for a transurethral resection of the bladder tumor, 18 were patients admitted for elective surgery for a non-malignant condition (benign prostate hyperplasia or upper urinary stone disease) and 17 were volunteering members of staff who provided two urine samples at least three months apart.

In each urine sample, a mean of 152 ± 128 OTUs were identified in total bacterial community. In each pair of samples from an individual subject, 41 ± 32 (27 ± 21%) of OTUs were overlapping, i.e. detected in both collections. These OTUs present in both collections covered in average 66.3 ± 29.4% (median 72.7%, IQR 49.7%) of the microbial community in urine samples in terms of relative abundance. There was a clear correlation between the number of overlapping OTUs and the relative abundance they were accounting for (Pearson’s R^2^ = 0.43, p < 0.001) (Fig 1).

**Fig 1:**
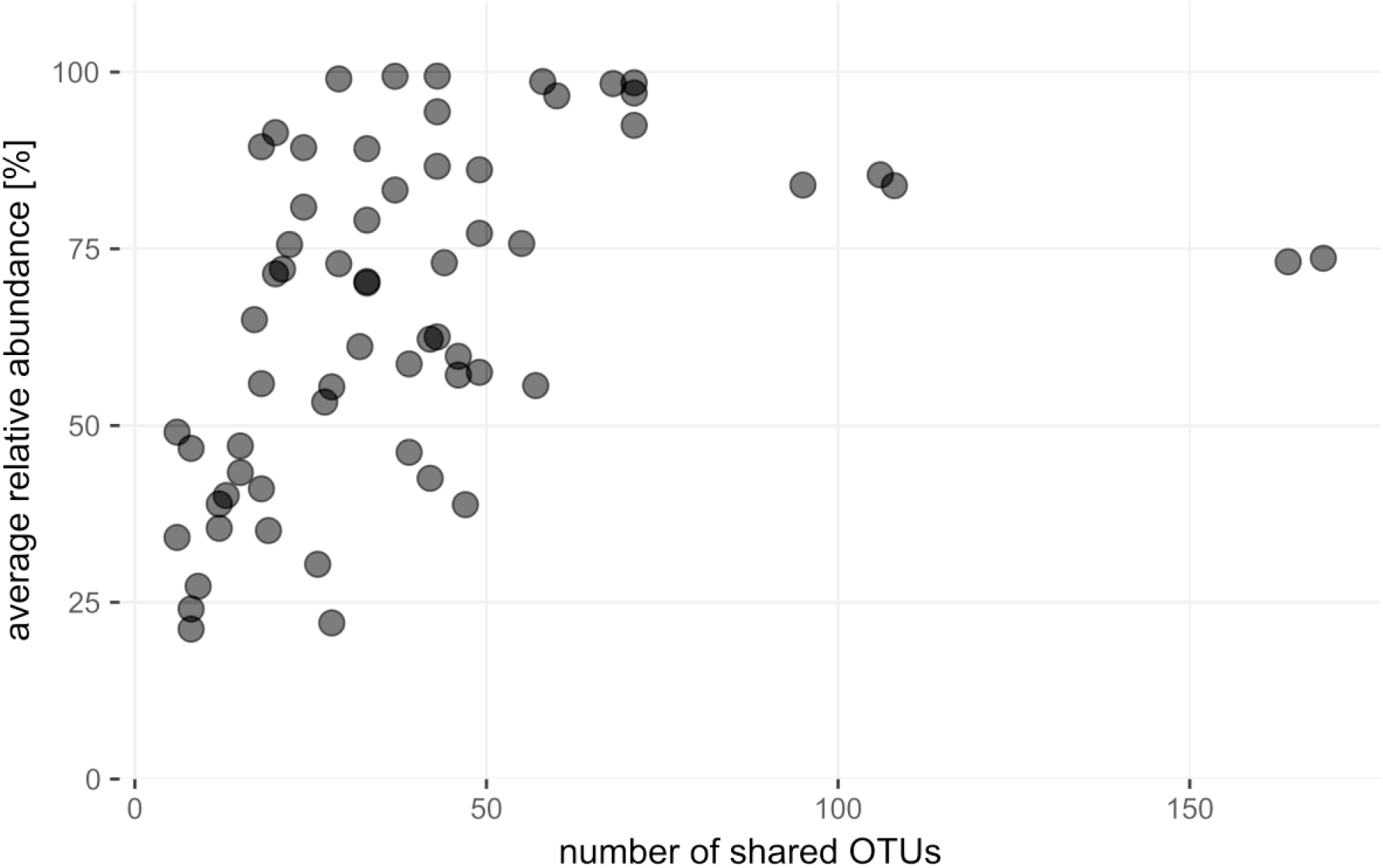
The correlation between the number of overlapping OTUs and the relative abundance they are covering. The more OTUs are shared between two samples of urine taken apart, the higher relative abundance they account for (Pearson’s R^2^ = 0.43, p<0.001).

Median value of Chao1 index was lower by 15.4 (IQR = 129.6) when comparing the second collection to the first one (Wilcoxon test, p < 0.05, Fig 2A). Shannon index showed no significant difference in community evenness between individual collections (Fig 2B). Comparison of groups with and without bladder cancer and groups with and without ureterolithiasis did not show differences in alpha diversity between these health conditions (S3 Table).

**Fig 2:**
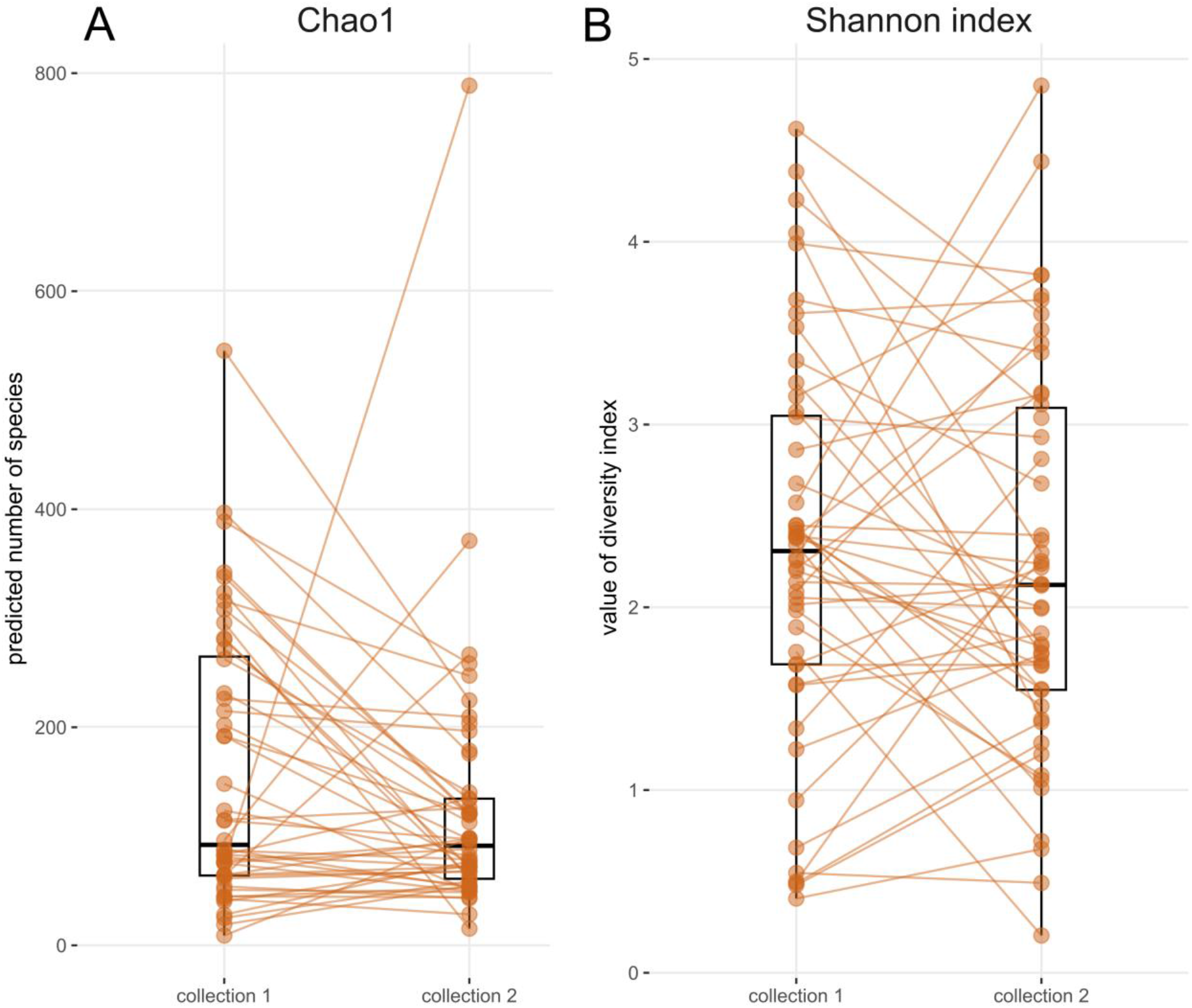
Microbial community indices of alpha diversity for two collection timepoints. (A) Chao1 and (B) Shannon index after 1,000× rarefaction. Boxplots show median, upper, and lower quartile, highest and lowest values. The first and second samples from the same subject are connected by orange lines.

In addition to alpha diversity metrics, the sum of bacterial relative abundance represented by overlapping taxa as well as their identity was evaluated in relation to biological sex. There was no difference in the number of overlapping OTUs and the relative abundance of overlapping community between males and females (Figs 3A, 3B) and between patients with bladder cancer and patients without cancer plus healthy volunteers (S1 Fig).

**Fig 3:**
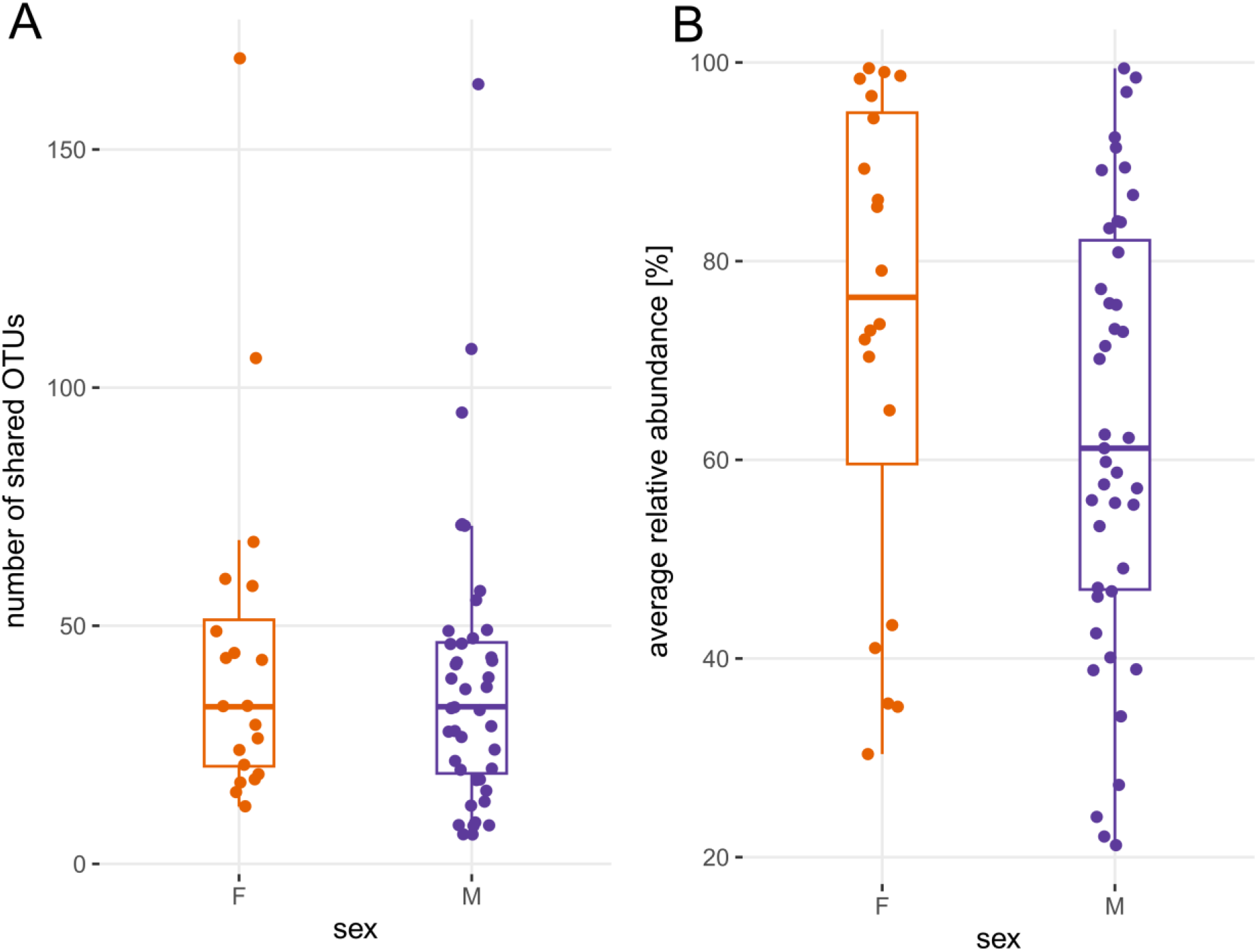
Overlap of the bacterial community from the two collection timepoints. (A) number of overlapping OTUs (e.g. taxa detected in both collections). (B) averaged sum of relative abundances of overlapping taxa from collections at two timepoints ([relative abundance of the overlapping OTUs at first sampling + relative abundance of the overlapping OTUs at second sampling] / 2). Boxplots show median, upper and lower quartile, highest and lowest values.

There was a non-significant trend towards lower overlap with increasing age of study subjects in both taxa count and relative abundance (Figs 4A and 4B). With a longer interval between the first and second urine collections the number of overlapping OTUs did not change significantly (Fig 5A). There was, however, a significantly decreased mean relative abundance of the overlapping taxa with time ([relative abundance of the overlapping OTUs at first sampling + relative abundance of the overlapping OTUs at second sampling] / 2; p < 0.05; Fig 5B).

**Fig 4:**
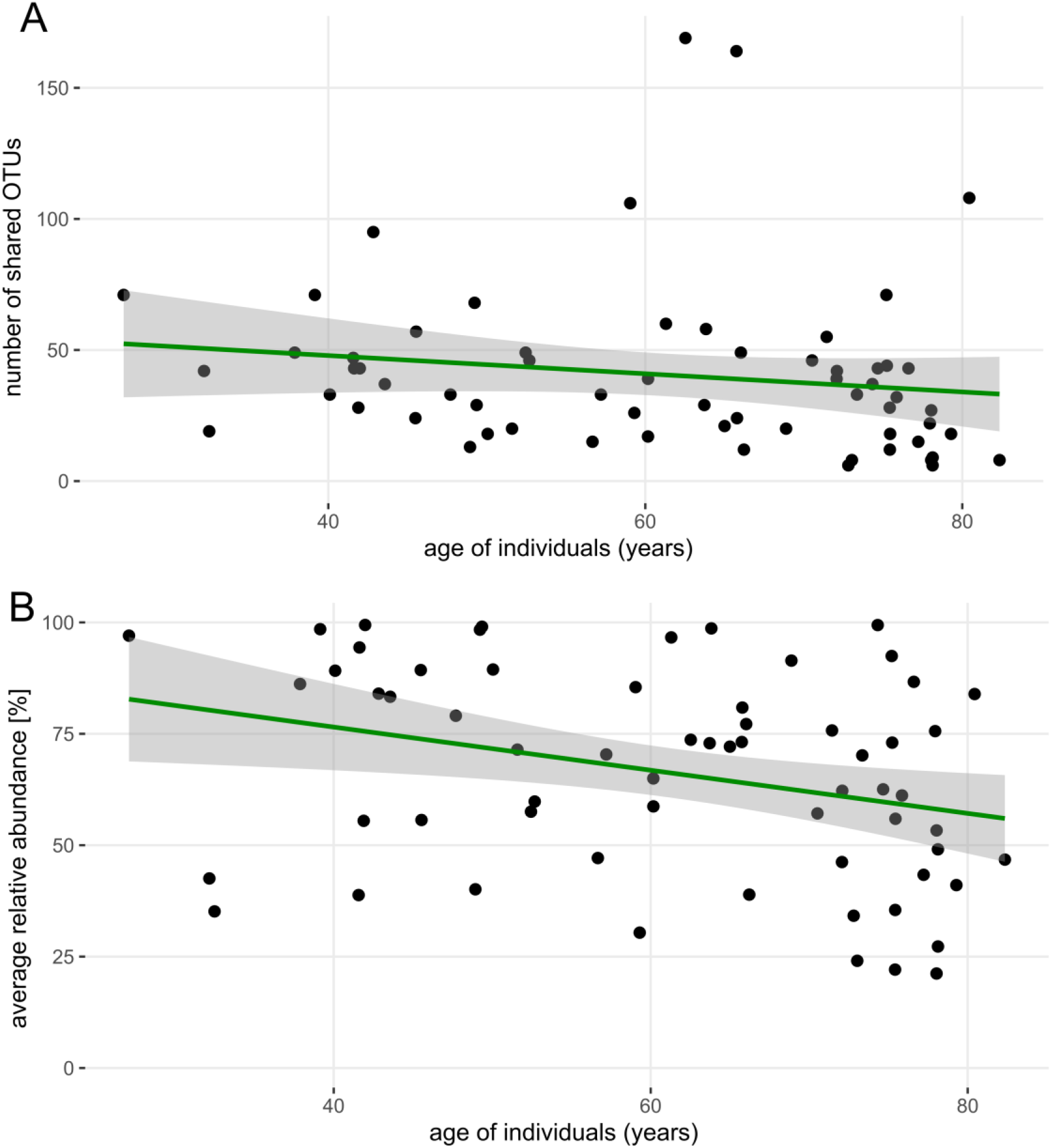
Relationship between taxa representation and age of subjects. (A) the number of OTUs which were detected in both collections as a function of age. (B) averaged sum of overlapping taxa relative abundance from collections at two timepoints as a function of age. Some urine pairs overlapping taxa represented nearly 100% of the community in both collections. In some cases, the relative abundance of the overlapping taxa was as low as 25%. Shaded area represents linear smoothing with 0.95 confidence interval.

**Fig 5:**
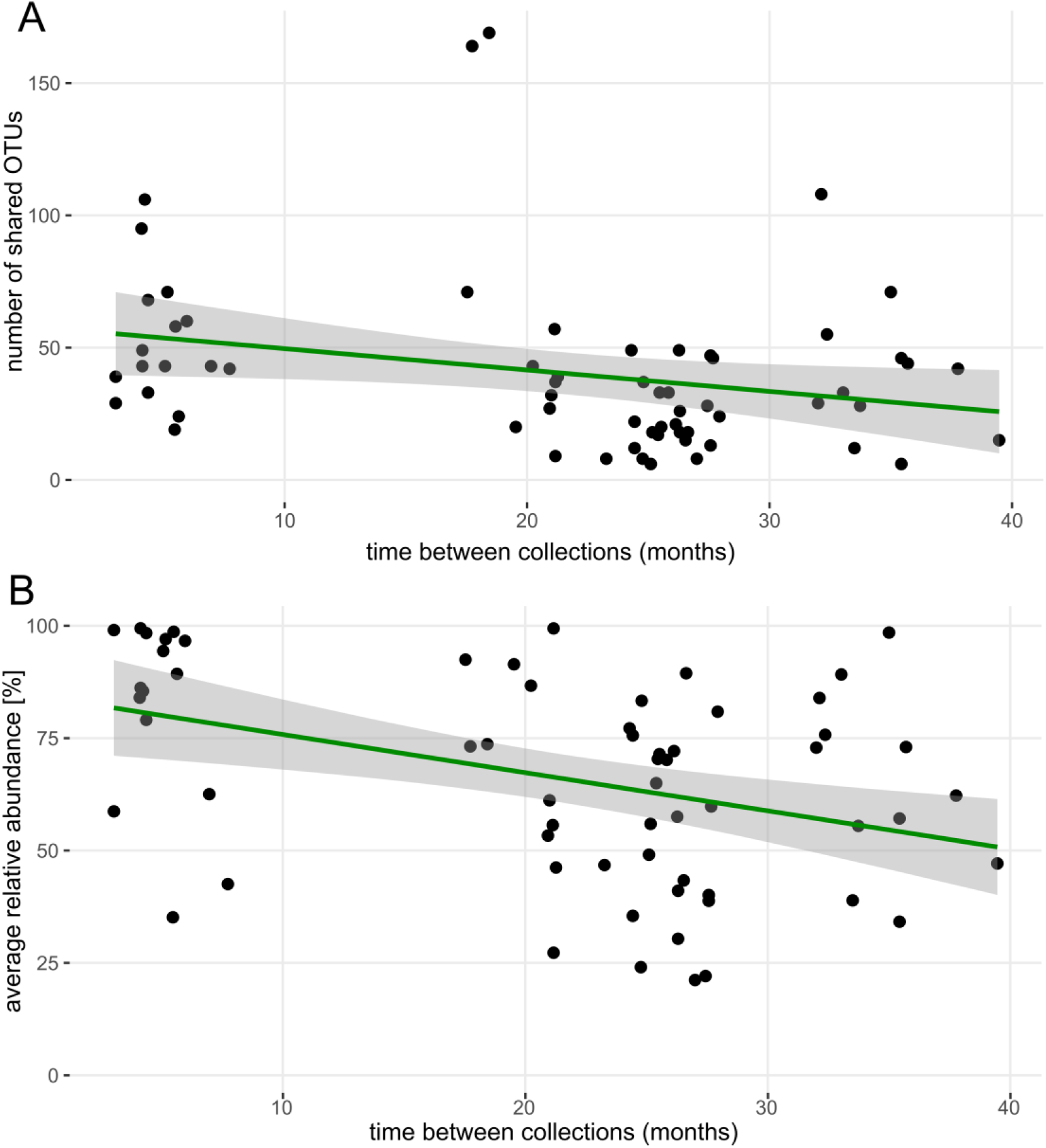
Relationship between overlapping bacterial community and time between collections. (A) number of OTUs which were detected in both collections as a function of time between collections. (B) averaged sum of overlapping taxa relative abundance as a function of time between collections (p < 0.05). Shaded area represents linear smoothing with 0.95 confidence interval.

In terms of beta diversity, the patients’ identity was a stronger driver for community composition than the time between collections (R^2^ = 0.04 and 0.02, respectively, p < 0.05). There were certain OTUs frequently observed in both paired samples and in more than 50% of male and female subjects (Table 1).

**Table 1.** Taxa detected in both the first and second urine sample by frequency.

| Genus (Class) | OTU ID | incidence in males | incidence in females |
| --- | --- | --- | --- |
|  |  | (%) | (%) |
| *Staphylococcus (Bacilli)* | CL00011 | 40 (93%) | 17 (85%) |
| unclass. *Enterobacteriaceae* |  |  |  |
| *(Gammaproteobacteria)* | CL00007 | 31 (72%) | 16 (80%) |
| *Cutibacterium (Actinobacteria)* | CL00024 | 34 (79%) | 14 (70%) |
| *Finegoldia (Clostridia)* | CL00021 | 29 (67%) | 14 (70%) |
| *Peptoniphilus (Clostridia)* | CL00035 | 27 (63%) | 14 (70%) |
| *Corynebacterium (Actinobacteria)* | CL00041 | 31 (72%) | 12 (60%) |
| unclass. *Enterobacteriaceae* |  |  |  |
| *(Gammaproteobacteria)* | CL00004 | 27 (63%) | 13 (65%) |
| *Streptococcus (Bacilli)* | CL00020 | 28 (65%) | 0 (0%) |
| *Fusobacterium (Fusobacteriia)* | CL00076 | 0 (0%) | 13 (65%) |
| *Enhydrobacter (Gammaproteobacteria)* | CL00050 | 26 (60%) | 0 (0%) |
| *Campylobacter (Campylobacteria)* | CL00013 | 0 (0%) | 12 (60%) |
| *Dialister (Negativicutes)* | CL00030 | 0 (0%) | 12 (60%) |
| *Gardnerella (Actinobacteria)* | CL00003 | 0 (0%) | 11 (55%) |
| *Bacteroidales (Bacteroidia)* | CL00006 | 0 (0%) | 11 (55%) |
| *Anaerococcus (Clostridia)* | CL00015 | 23 (53%) | 0 (0%) |
| *Corynebacterium (Actinobacteria)* | CL00054 | 22 (51%) | 0 (0%) |

Taxa detected in both the first and second urine sample with more than 50% prevalence in either
sex.

## Discussion

Since the discovery of the human UM in 2011 (1,2), it has been associated with various pathological conditions. Twelve years later, there is not only a lack of description of normal UM, but also a lack of deeper understanding of UM dynamics. However, this knowledge is crucial to draw conclusions about the role of UM in the onset and development of disease. For example Isali et al. (27) recently pointed out that it is difficult to characterize causality in microbiome research of bladder cancer and even with known composition of microbial taxa, these results cannot be readily translated to therapeutic applications. This study aims to uncover the effects of time on the UM composition using repeated sampling of study participants. Furthermore, we want to characterize the overlap of the microbial community from the same subject over a period of several months.

Microbial communities of other human body niches are known to be stable over time. (6–8) For the urinary tract, there has been little evidence on this topic. The first endeavor investigating UM short-term stability on 14 subjects of both sexes and various ages using 16S rRNA gene sequencing found that the voided urinary microbiota composition remained stable on a short-term basis (several days apart). (28) Eight women were observed longitudinally for three months with daily culture of mid-stream urine (MSU) and periurethral swab; UM was undergoing fluctuations associated with menstruation and sexual activity. (29) In a cohort of 10 women who provided mid-stream urine samples 2.5 years apart, high intraindividual variability was observed with only 29% (range 9–42%) of species overlapping from the first and second sample even though the sampling was performed uniformly in the third week of the menstrual cycle. Of note, the authors chose to use an expanded quantitative urine culture (EQUC) protocol instead of 16S rRNA gene sequencing to assess the microbial diversity. (30) In 13 pregnant women whose mid-stream urine samples were collected in the first and second trimester and subjected to 16S rRNA gene sequencing, the UM was stable with *Lactobacillus* being the dominant microorganism in the majority of women. (31)

In the present study, we show that the number of bacterial taxa overlapping in two urine collections usually does not exceed 50 (Figs 1, 3B, 4A, 5A) and that the relative abundance of these overlapping OTUs averages between 60% and 70% sometimes even reaching 90–100% (Figs 4B and 5B). There was no significant difference between men and women in the number of shared OTUs or in the relative abundance represented by these OTUs (Figs 3A, 3B). The bladder cancer status of patients also had no effect on the number of shared OTUs or their sum of abundance (S1 Fig).

Using the Chao1 estimator, we have shown that the richness of the microbial community changes between collections, i.e. there is some variation in the number of resident bacteria in urine samples taken months apart (Fig 2A). The relative abundance of overlapping OTUs may vary considerably (Fig 2B) and this phenomenon becomes more significant as the time interval between collections increases (Fig 5B).

At the community composition level (beta diversity), greater variability was explained by the subject identity than by the effect of collection time. Therefore, the differences in community composition within a subject were smaller than the differences between individuals.

Based on the design of our previous project (10), to which the present study is an extension, 26 of 63 (41%) urine pairs contained catheterized urine from the first collection (all second samples were MSU). When MSU was compared to MSU in both samples, there were 47 ± 32 overlapping OTUs; when catheterized urine from the first collection was compared to MSU from the second sampling, the proportion of overlapping OTUs decreased to 32 ± 31 OTUs. This difference was statistically significant (p = 0.01) and is consistent with previous research reporting microbial communities in catheterized samples differ from MSU. (12)

We have noted that certain OTUs were observed in both paired samples in more than 50% of subjects (Table 1). These might represent candidate OTUs in the quest for a core urinary microbiota. Whether such phenomenon exists is still a matter of debate but of the taxa listed in Table 1, the following have been consistently reported in previous urinary microbiota studies (in alphabetical order): *Corynebacterium, Finegoldia, Gardnerella, Peptoniphilus, Staphylococcus* and *Streptococcus* (1,32–35). Aerobic subgroup of these taxa (like *Corynebacterium, Staphylococcus* and *Streptococcus* are also present in a recent genome collection of bladder-specific isolates and thus their metabolic potential might be assessed. (36) Other taxa like *Peptoniphilus* are more elusive due to their anaerobic growth. The fact that *Lactobacillus* does not feature among the top OTUs in our dataset may be explained by the postmenopausal status of the study female participants (37); indeed, *Lactobacillus* was detected in both paired samples in only 40% of women.

We analyzed the dynamics of the microbial community in individual sample pairs (S2 Fig) to explain very low community overlap in some cases. In some of the pairs with the lowest overlap, a particular OTU was present with a high relative abundance (>50%) in one sample and absent in the other. These OTUs were identified as (in alphabetical order) *Arcanobacterium, Enterobacter/Morganella, Idiomarina, Lactobacillus, Pseudomonas and Ureaplasma*.

Some methodological weaknesses of our study deserve attention: 1) Urine pairs collected at shorter time intervals between collections were from younger subjects, while paired samples from older participants were separated by a longer time gap. This may have led to a systematic bias favoring the statistical significance of the trend seen in Fig 5B; 2) while none of the study subjects had taken antibiotics six weeks prior to the first urine collection, the absence of antibiotic treatment was less rigorously documented at the second urine sampling. Therefore, we cannot rule out the possibility that a shift in the richness or diversity of the UM was due to undocumented antibiotic exposure; 3) patients who harbored urinary stones at first sampling may have been stone-free at second sampling. While these shortcomings weaken the statistical significance of some of our results, they strengthen the claim that UM undergoes only limited change in time.

## Ethical approval

The study was approved by the Ethics Committee of the Institute for Clinical and Experimental Medicine and Thomayer Hospital in March 2019 (docket no. G-19-01) and extension of the study was approved in May 2021 (docket no. 125699/21).

## Author Contributions

JH, VH, and RZ designed the study. JH, VH, PB, PC, and RZ performed sample collection. PB and PC performed lab research. VT and PTD analyzed the data. JH assisted with data interpretation. JH and VT wrote the manuscript.

---

**S1 Fig: Overlap of the bacterial community from the two collection timepoints**. (A) number of overlapping OTUs (e.g. taxa detected in both collections). (B) averaged sum of relative abundances of overlapping taxa from collections at two timepoints ([relative abundance of the overlapping OTUs at first sampling + relative abundance of the overlapping OTUs at second sampling] / 2). Boxplots show median, upper and lower quartile, highest and lowest values for groups of patients with and without bladder cancer.

**S2 Fig: The dynamics of the bacterial community shared among individual sample pairs**. Individual facets represent patients with two collections each, numbers in facet headers represent patient IDs. Shared OTUs are depicted as points and direction of change in their relative abundance between collections is visualized by connecting lines.

**S1 Table: The STORMS checklist**. Form was downloaded from https://stormsmicrobiome.org.

**S2 Table: Metadata and individual data points from analysis of 69 subjects which were recruited for the study**. CKD-EPI eGFR_ml_min - Glomerular Filtration Rate, RT_MP_in_anam – pelvic radiotherapy in patient history.an

**S3 Table: Values of alpha diversity for selected groups of subjects**. Average value ± standard deviation is given; upper indices mark pairs with significant differences.

