## Supplementary Table 1 for "Short- and mid-term temporal variability of the human urinary microbiota: an observational cohort study"

Supplementary Table 1 - The STORMS checklist, form was downloaded from [https://stormsmicrobiome.org](https://stormsmicrobiome.org/).

| Number | Item | Recommendation | Item Source | Additional Guidance | Yes/No/NA | Comments or location in manuscript |
| --- | --- | --- | --- | --- | --- | --- |
| **Abstract** | | | | | | |
| 1.0 | Structured or Unstructured Abstract | Abstract should include information on background, methods, results, and conclusions in structured or unstructured format. | STORMS |  | Yes |  |
| 1.1 | Study Design | State study design in abstract. | STORMS | See 3.0 for additional information on study design. | Yes |  |
| 1.2 | Sequencing methods | State the strategy used for metagenomic classification. | STORMS | For example, targeted 16S by qPCR or sequencing, shotgun metagenomics, metatranscriptomics, etc. | Yes | Methods |
| 1.3 | Specimens | Describe body site(s) studied. | STORMS |  | Yes | Methods |
| **Introduction** | | | | | | |
| 2.0 | Background and Rationale | Summarize the underlying background, scientific evidence, or theory driving the current hypothesis as well as the study objectives. | STORMS |  | Yes | Introduction |
| 2.1 | Hypotheses | State the pre-specified hypothesis. If the study is exploratory, state any pre-specified study objectives. | STORMS | The study is exploratory. The objective was to assess the temporal variation in the urinary microbiota composition. | Yes | Introduction |
| **Methods** | | | | | | |
| 3.0 | Study Design | Describe the study design. | STORMS | [Observational (Case-Control, Cohort, Cross-sectional survey, etc.) or Experimental (Randomized controlled trial, Non-randomized controlled trial, etc.). For a brief description of common study designs see: DOI: 10.11613/BM.2014.022](https://doi.org/10.1592/phco.30.10.973)  [If applicable, describe any blinding (e.g. single or double-blinding) used in the course of the study.](https://doi.org/10.1592/phco.30.10.973) | Yes | Methods |
| 3.1 | Participants | State what the population of interest is, and the method by which participants are sampled from that population. Include relevant information on physiological state of the subjects or stage in the life history of disease under study when participants were sampled. | STORMS | The population consists of adults who attended the hospital for an elective endoscopic procedure either for bladder cancer or treatment for urinary stone disease/benign prostate hyperplasia. The study cohort was expanded using healthy volunteering hospital staff. | Yes | Methods, PRJNA1093489 |
| 3.2 | Geographic location | State the geographic region(s) where participants were sampled from. | MIxS: geographic location (country and/or sea,region) | Geographic coordinates can be reported to prevent potential ambiguities if necessary. | Yes | Methods |
| 3.3 | Relevant Dates | State the start and end dates for recruitment, follow-up, and data collection. | STORMS | Recruitment is the period in which participants are recruited for the study. In longitudinal studies, follow-up is the date range in which participants are asked to complete a specific assessment. Finally, data collection is the total period in which data is being collected from participants including during initial recruitment through all follow-ups. | Yes | Methods |
| 3.4 | Eligibility criteria | List any criteria for inclusion and exclusion of recruited participants. | Modified STROBE | Among potential recruited participants, how were some chosen and others not? This could include criteria such as sex, diet, age, health status, or BMI.  If there is a primary and validation sample, describe inclusion/exclusion criteria for each. | Yes | Methods |
| 3.5 | Antibiotics Usage | List what is known about antibiotics usage before or during sample collection. | STORMS | If participants were excluded due to current or recent antibiotics usage, state this here.  Other factors (e.g. proton pump inhibitors, probiotics, etc.) that may influence the microbiome should also be described as well. | Yes | Methods |
| 3.6 | Analytic sample size | Explain how the final analytic sample size was calculated, including the number of cases and controls if relevant, and reasons for dropout at each stage of the study. This should include the number of individuals in whom microbiome sequencing was attempted and the number in whom microbiome sequencing was successful. | STORMS | Consider use of a flow diagram (see template at https://stormsmicrobiome.org/figures). Also state sample size in abstract.  If power analysis was used to calculate sample size, describe those calculations. | Yes | Methods |
| 3.7 | Longitudinal Studies | For longitudinal studies, state how many follow-ups were conducted, describe sample size at follow-up by group or condition, and discuss any loss to follow-up. | STORMS | If there is loss to follow-up, discuss the likelihood that drop-out is associated with exposures, treatments, or outcomes of interest. | Yes | Methods |
| 3.8 | Matching | For matched studies, give matching criteria. | Modified STROBE | "Matched" refers to matching between comparable study participants as cases and controls or exposed / unexposed.  Indicate whether participants were individual or frequency matched and in what ratio were they matched (e.g. 1 case to 1 control). | NA |  |
| 3.9 | Ethics | State the name of the institutional review board that approved the study and protocols, protocol number and date of approval, and procedures for obtaining informed consent from participants. | STORMS | Ethics Committee of the Institute for Clinical and Experimental Medicine and Thomayer Hospital with Multi-center Competence | Yes | Methods |
| 4.0 | Laboratory methods | State the laboratory/center where laboratory work was done. | STORMS | Provide a reference to complete lab protocols if previously published elsewhere such as on protocols.io. Note any modifications of lab protocols and the reason for protocol modifications. | Yes | Methods |
| 4.1 | Specimen collection | State the body site(s) sampled from and how specimens were collected. | MIxS: sample collection device or method; host body site | Use terms from the Uber-anatomy Ontology (https://www.ebi.ac.uk/ols/ontologies/uberon) to describe body sites in a standardized format. | Yes | Methods |
| 4.2 | Shipping | Describe how samples were stored and shipped to the laboratory. | STORMS | Include length of time from collection to receipt by the lab and if temperature control was used during shipping. | Yes | Methods |
| 4.3 | Storage | Describe how the laboratory stored samples, including time between collection and storage and any preservation buffers or refrigeration used. | STORMS | State where each procedure or lot of samples was done if not all in the same place.  Include reagent/lot/catalogue #s for storage buffers. | Yes | Methods |
| 4.4 | DNA extraction | Provide DNA extraction method, including kit and version if relevant. | MIxS: nucleic acid extraction | If any DNA quantification methods were used prior to DNA amplification or at the pooling step of library preparation, state so here. | Yes | Methods, quantification before library prep using Qubit |
| 4.5 | Human DNA sequence depletion or microbial DNA enrichment | Describe whether human DNA sequence depletion or enrichment of microbial or viral DNA was performed. | STORMS |  | Yes | Methods |
| 4.6 | Primer selection | Provide primer selection and DNA amplification methods as well as variable region sequenced (if applicable). | MIxS: pcr primers |  | Yes | Methods |
| 4.7 | Positive Controls | Describe any positive controls (mock communities) if used. | STORMS | If used, should be deposited under guidance provided in the 8.X items. | NA |  |
| 4.8 | Negative Controls | Describe any negative controls if used. | STORMS | If used, should be deposited under guidance provided in the 8.X items. | Yes | Methods |
| 4.9 | Contaminant mitigation and identification | Provide any laboratory or computational methods used to control for or identify microbiome contamination from the environment, reagents, or laboratory. | STORMS | Includes filtering of reagents and other steps to minimize contamination. It is relevant to state whether the specimens of interest have low microbial load, which makes contamination especially relevant. | Yes | Methods |
| 4.10 | Replication | Describe any biological or technical replicates included in the sequencing, including which steps were replicated between them. | STORMS | Replication may be biological (redundant biological specimens) or technical (aliquots taken at different stages of analysis) and used in extraction, sequencing, preprocessing, and/or data analysis. | Yes | Methods |
| 4.11 | Sequencing strategy | Major divisions of strategy, such as shotgun or amplicon sequencing. | MIxS: sequencing method | For amplicon sequencing (for example, 16S variable region), state the region selected. State the model of sequencer used. | Yes | Methods |
| 4.12 | Sequencing methods | State whether experimental quantification was used (QMP/cell count based, spike-in based) or whether relative abundance methods were applied. | STORMS | These include read length, sequencing depth per sample (average and minimum), whether reads are paired, and other parameters. | Yes | Methods |
| 4.13 | Batch effects | Detail any blocking or randomization used in study design to avoid confounding of batches with exposures or outcomes. Discuss any likely sources of batch effects, if known. | STORMS | Sources of batch effects include sample collection, storage, library preparation, and sequencing and are commonly unavoidable in all but the smallest of studies. | Yes | Methods |
| 4.14 | Metatranscriptomics | Detail whether any mRNA enrichment was performed and whether/how retrotranscription was performed prior to sequencing. Provide size range of isolated transcripts. Describe whether the sequencing library was stranded or not. Provide details on sequencing methods and platforms. | STORMS | Provide details on any internal standards which may have been used as well as parameters and versions of any software or databases used. | N/A |  |
| 4.15 | Metaproteomics | Detail which protease was used for digestion. Provide details on proteomic methods and platforms (e.g. LC-MS/MS, instrument type, column type, mass range, resolution, scan speed, maximum injection time, isolation window, normalised collision energy, and resolution). | STORMS | Provide details on any internal standards which may have been used as well as parameters and versions of any software or databases used. | N/A |  |
| 4.16 | Metabolomics | Specify the analytic method used (such as nuclear magnetic resonance spectroscopy or mass spectrometry). For mass spectrometry, detail which fractions were obtained (polar and/or non-polar) and how these were analyzed. Provide details on metabolomics methods and platforms (e.g. derivatization, instrument type, injection type, column type and instrument settings). | STORMS | Provide details on any internal standards which may have been used as well as parameters and versions of any software or databases used. | N/A |  |
| 5.0 | Data sources/  measurement | For each non-microbiome variable, including the health condition, intervention, or other variable of interest, state how it was defined, how it was measured or collected, and any transformations applied to the variable prior to analysis. | MIxS: host disease status | State any sources of potential bias in measurements, for example multiple interviewers or measurement instruments, and whether these potential biases were assessed or accounted for in study design.  Use terms from a standardized ontology such as the Experimental Factor Ontology (https://www.ebi.ac.uk/efo/) to describe variables of interest in a standardized format. | Yes | PRJNA1093489 |
| 6.0 | Research design for causal inference | Discuss any potential for confounding by variables that may influence both the outcome and exposure of interest. State any variables controlled for and the rationale for controlling for them. | STORMS | For causal inference, this item refers to describing the assumptions that would be required to draw causal inferences from observational data. See Vujkovic-Cvijin, I., Sklar, J., Jiang, L. et al. Host variables confound gut microbiota studies of human disease. Nature 587, 448–454 (2020). https://doi.org/10.1038/s41586-020-2881-9 for more details on confounding in observational microbiome studies.  For example, hypothesized confounders may be controlled for by multivariable adjustment. Consider using a directed acyclic graph (DAG) to describe your causal model and justify any variables controlled for. DAGs can be made using [www.dagitty.net](http://www.dagitty.net/). | Yes | Discusssion |
| 6.1 | Selection bias | Discuss potential for selection or survival bias. | STORMS | Selection bias can occur when some members of the target study population are more likely to be included in the study/final analytic sample than others. Some examples include survival bias (where part of the target study population is more likely to die before they can be studied), convenience sampling (where members of the target study population are not selected at random), and loss to follow-up (when probability of dropping out is related to one of the things being studied). | Yes | Despite the random selection of study subjects, not all groups have random distribution. |
| 7.0 | Bioinformatic and Statistical Methods | Describe any transformations to quantitative variables used in analyses (e.g. use of percentages instead of counts, normalization, rarefaction, categorization). | STORMS | If a variable is analyzed using different transformations, state rationale for the transformation and for each analyses which version of the variable is used.  In case of any complex or multistep transformations, give enumerated instructions for reproducing those transformations. | Yes | Methods |
| 7.1 | Quality Control | Describe any methods to identify or filter low quality reads or samples. | MIxS: sequence quality check | If samples were excluded based on quality or read depth, list the criteria used, the number of samples excluded, and the final sample size after quality control. | Yes | Methods |
| 7.2 | Sequence analysis | Describe any taxonomic, functional profiling, or other sequence analysis performed. | MIxS: feature prediction; similarity search method |  | Yes | Methods |
| 7.3 | Statistical methods | Describe all statistical methods. | Modified STROBE | Describe any statistical tests used, exploratory data analysis performed, dimension reduction methods/unsupervised analysis, alpha/beta metrics, and/or methods for adjusting for measurement bias.  If multiple statistical methods are possible, discuss why the methods used were selected.  If a multiple hypothesis testing correction method was used, describe the type of correction used.  State which taxonomic levels are analyzed. | Yes | Methods |
| 7.4 | Longitudinal analysis | If the study is longitudinal, include a section that explicitly states what analysis methods were used (if any) to account for grouping of measurements by individual or patterns over time. | STORMS |  | Yes | Averaged abundances used for collections from the same patient |
| 7.5 | Subgroup analysis | Describe any methods used to examine subgroups and interactions. | STROBE |  | Yes | PERMANOVA, Methods |
| 7.6 | Missing data | Explain how missing data were addressed. | STROBE | "Missing data" refers to participant measurements such as covariates, exposures, outcomes, or time points that should have been collected but were not, not to zeros in taxonomic abundance tables or data points not applicable to that observation. | NA |  |
| 7.7 | Sensitivity analyses | Describe any sensitivity analyses. | STROBE |  | NA |  |
| 7.8 | Findings | State criteria used to select findings for reporting. | STORMS | For example, false discovery rate with total number of tests, effect size threshold, significance threshold, microbes of interest. | Yes | significance threshold, microbes of interest |
| 7.9 | Software | Cite all software (including read mapping software) and databases (including any used for taxonomic reference or annotating amplicons, if applicable) used. Include version numbers. | Modified STREGA | Installed packages, add-ons or libraries should be stated and cited in addition to the software used.  All parameters employed that differ from the default of that software/version should be provided.  This is in addition to, not a replacement for, publishing of code as outlined in the section Reproducible Research. | Yes | Methods |
| 8.0 | Reproducible research | Make a statement about whether and how others can reproduce the reported analysis. | STORMS | Any protected information that has been excluded or provided under controlled access should be listed along with any relevant data access procedures. "On request from authors" is not sufficiently detailed; formal data access procedures and conditions should be defined.  If data are unavailable, state so clearly.  Consider using a specialized rubric for reproducible research (such as:<https://mbio.asm.org/content/9/3/e00525-18.short)>.  Consider preregistering the study protocol (such as o[n osf.](http://osf.io/)io or<https://plos.org/open-science/preregistration/).> | Yes | PRJNA1093489 |
| 8.1 | Raw data access | State where raw data may be accessed including demultiplexing information. | STORMS | Robust, long-term databases such as those hosted by NCBI and EBI are preferred. If using a private repository, provide rationale. | Yes | PRJNA1093489 |
| 8.2 | Processed data access | State where processed data may be accessed. | STORMS | Unfiltered data should be provided.  Robust, long-term databases such as those hosted by NCBI and EBI-EMBL are preferred. Repositories like zenodo (https://zenodo.org/) or publisso (https://www.publisso.de/en/working-for-you/doi-service/)  can be used to provide a DOI and long-term storage for processed datasets, even those which cannot be published openly. | Yes | 10.6084/m9.figshare.26046355 |
| 8.3 | Participant data access | State where individual participant data such as demographics and other covariates may be accessed, and how they can be matched to the microbiome data. | STORMS | If re-categorized, transformed, or otherwise derived variables were used in the analysis, these variables or code for deriving them should be provided.  Examples of how participant data can be matched to microbiome data are: using the same set of anonymized identifiers, or using different anonymized identifiers but providing a map.  Provided data should be sufficient to independently replicate the current analysis. | Yes | PRJNA1093489 |
| 8.4 | Source code access | State where code may be accessed. | STORMS | If a standard or formalized workflow was employed, reference it here. | Yes | Methods, prhttps://github.com/TlaskalV/human-urinary-microbiome |
| 8.5 | Full results | Provide full results of all analyses, in computer-readable format, in supplementary materials. | STORMS | For example, any fold-changes, p-values, or FDR values calculated, provided as a spreadsheet.  Use a machine-readable, plain-text format such as csv or tsv. | Yes | 10.6084/m9.figshare.26046355 |
| **Results** | | | | | | |
| 9.0 | Descriptive data | Give characteristics of study participants (e.g. dietary, demographic, clinical, social) and information on exposures and potential confounders. | STROBE | Typically reported in a table included in the paper or as a supplementary table. Indicate number of participants with missing data for each variable of interest.  This includes environmental and lifestyle factors that may affect the relationship between the microbiome and the condition of interest. Participant diet and medication use should be summarized, if known.  At minimum, age and sex of all participants should be summarized. | Yes | PRJNA1093489 |
| 10.0 | Microbiome data | Report descriptive findings for microbiome analyses with all applicable outcomes and covariates. | STORMS | This includes measures of diversity as well as relative abundances. These descriptive findings should be reported both for the sample overall and for individual groups. | Yes | Results |
| 10.1 | Taxonomy | Identify taxonomy using standardized taxon classifications that are sufficient to uniquely identify taxa. | STORMS | If not using full taxonomic hierarchy, make sure it is clear whether names stated are species, genera, family, etc.  Italicize genus/species pairs. Consult journal guidelines or standardized references on taxonomic nomenclature. For instance,<https://wwwnc.cdc.gov/eid/page/scientific-nomenclature> | Yes | Results |
| 10.2 | Differential abundance | Report results of differential abundance analysis by the variable of interest and (if applicable) by time, clearly indicating the direction of change and total number of taxa tested. | STORMS | If there are more than two groups, include omnibus (multigroup) test results if applicable to the research question.  If applicable, reported effect sizes should include a measure of uncertainty such as the confidence interval. | Yes | Results |
| 10.3 | Other data types | Report other data analyzed--e.g. metabolic function, functional potential, MAG assembly, and RNAseq. | STORMS |  | N/A |  |
| 10.4 | Other statistical analysis | Report any statistical data analysis not covered above. | STORMS | This could include subgroup analysis, sensitivity analyses, and cluster analysis.  Visualizations should be easily interpretable and colorblind-friendly. The caption and/or main text should provide a detailed description of visualizations for visually-impaired readers. | NA |  |
| **Discussion** | | | | | | |
| 11.0 | Key results | Summarise key results with reference to study objectives | STROBE |  | Yes | Discussion |
| 12.0 | Interpretation | Give a cautious overall interpretation of results considering objectives, limitations, multiplicity of analyses, results from similar studies, and other relevant evidence. | STROBE | Define or clarify any subjective terms such as "dominant," "dysbiosis," and similar words used in interpretation of results.  When interpreting the findings, consider how the interpretation of the findings may be summarized or quoted for the general public such as in press releases or news articles.  If causal language is used in the interpretation (such as "alters," "affects," "results in," "causes," or "impacts"), assumptions made for causal inference should be explicitly stated as part of 6.0 and 13.0.  Distinguish between function potential (ie inferred from metagenomics) and observed activity (ie metatranscriptomic, metabolomic, proteomic) if discussing microbial function. | Yes | Discussion |
| 13.0 | Limitations | Discuss limitations of the study, taking into account sources of potential bias or imprecision. | STROBE | Also consider limitations resulting from the methods (especially novel methods), the study design, and the sample size. | Yes | Discussion |
| 13.1 | Bias | Discuss any potential for bias to influence study findings. | STORMS | May include sampling method, representativeness of study participants, or potential confounding. | Yes | Discussion |
| 13.2 | Generalizability | Discuss the generalisability (external validity) of the study results | STROBE | To what populations or other settings do you expect the conclusions to generalize? | Yes | Discussion |
| 14.0 | Ongoing/future work | Describe potential future research or ongoing research based on the study's findings. | STORMS |  | Yes | Discussion |
| **Other information** | | | | | | |
| 15.0 | Funding | Give the source of funding and the role of the funders for the present study and, if applicable, for the original study on which the present article is based | STROBE |  | Yes |  |
| 15.1 | Acknowledgements | Include acknowledgements of those who contributed to the research but did not meet critera for authorship. | STORMS | For general guidelines on authorship, see [http://www.icmje.org](http://www.icmje.org/) and<https://www.elsevier.com/authors/journal-authors/policies-and-ethics/credit-author-statement> | Yes |  |
| 15.2 | Conflicts of Interest | Include a conflicts of interest statement. | STORMS |  | Yes |  |
| 16.0 | Supplements | Indicate where supplements may be accessed and what materials they contain. | STORMS |  | Yes |  |
| 17.0 | Supplementary data | Provide supplementary data files of results with for all taxa and all outcome variables analyzed. Indicate the taxonomic level of all taxa. | STORMS | Depending on the analysis performed, examples of the supplemental results included could be mean relative abundance, differential abundance, raw p-value, multiple hypothesis testing-adjusted p-values, and standard error.  All discussed taxa should include the taxonomic level (e.g. class, order, genus). | Yes |  |
