## Supplementary figures and images for "Short- and mid-term temporal variability of the human urinary microbiota: an observational cohort study"

### Supplementary Figure 1

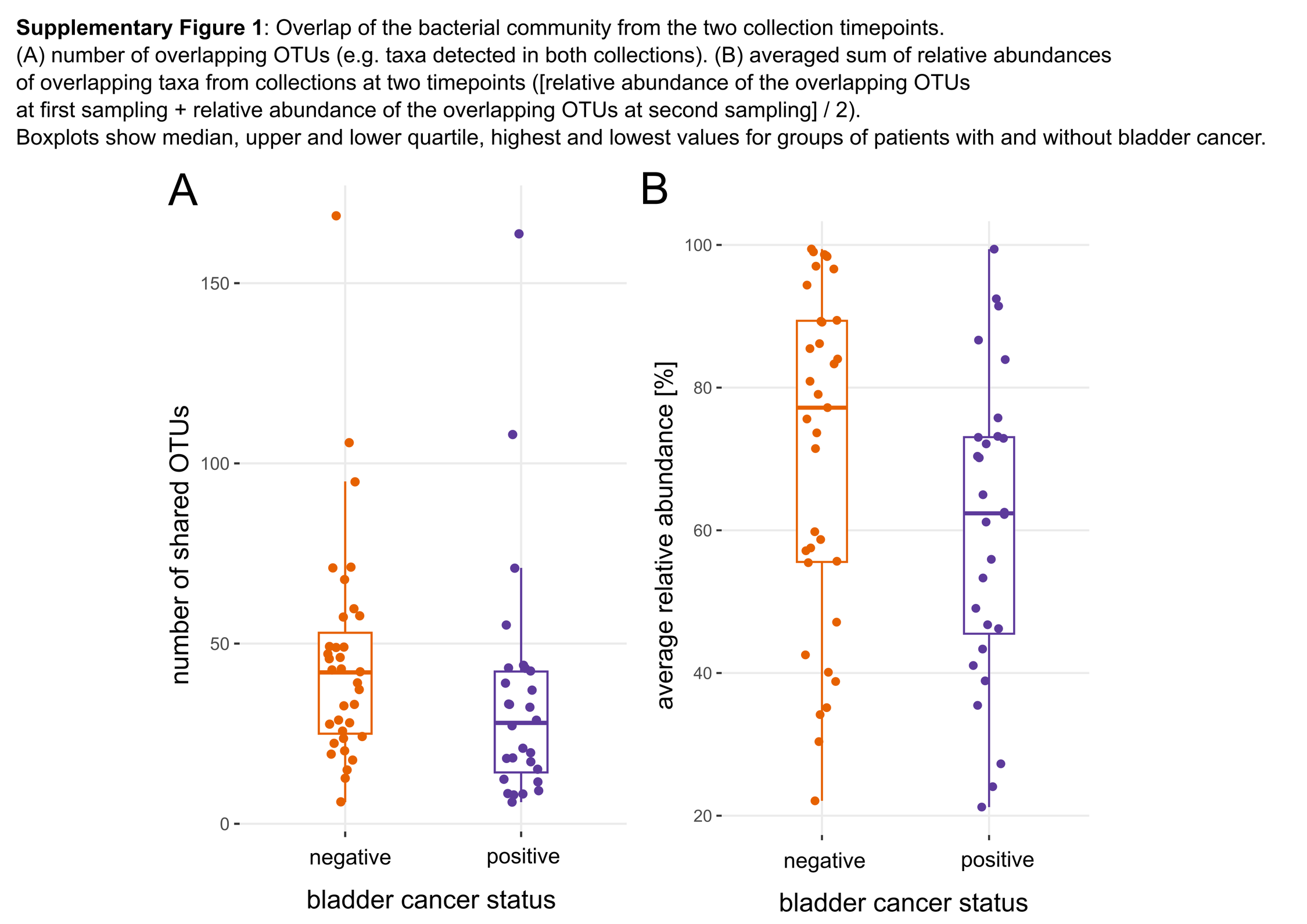

### Supplementary Figure 2

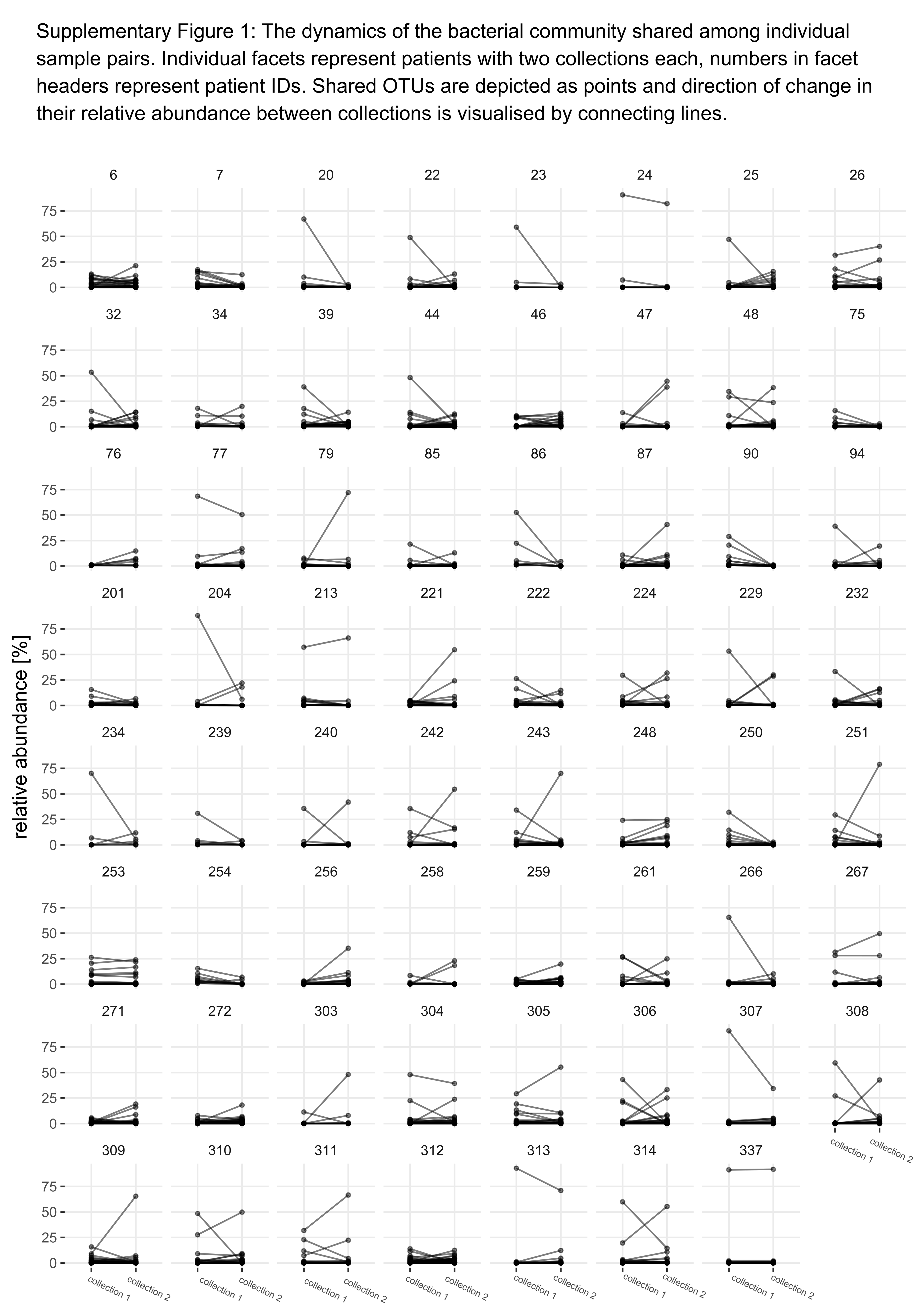
